# Application of Surgical Lead Management and Reconfigurable Coil Technology to Reduce RF Heating of DBS Implants during MRI at 3T Under Variant Body Compositions

**DOI:** 10.1101/2020.06.25.170159

**Authors:** Bhumi Bhusal, Behzad Elahi, Boris Keil, Joshua Rosenow, Ehsan Kazemivalipour, Laleh Golestanirad

**Author notes:** Corresponding author: L Golestanirad is with Department of Radiology and Department of Biomedical Engineering, Northwestern University, Chicago, IL, 60611 USA.

## Abstract

Patients with active implants such as deep brain stimulation (DBS) devices, have limited access to magnetic resonance imaging (MRI) due to risks of RF heating. With an aging population, the prevalence of neurodegenerative and vascular disease increases; and so does the indication for MRI exams in patients with such implants. In response to this growing need for MRI, many groups have investigated strategies to mitigate the RF heating of the implants. These efforts, however, have relied either on simulations with homogenous body models or simplified phantom experiments (box shaped phantom with single tissue). It is well established, however, that the shape and heterogeneity of human body affects the distribution of MRI electric fields, which by proxy, alters the RF heating of an implant inside the body. In this contribution, we applied numerical simulations and phantom experiments to examine the effectiveness of RF heating mitigation strategies under variant patient body compositions, focusing on two recently proposed techniques: (a) surgical modification of DBS lead trajectories inside the body, and (b) use of a patient-adjustable reconfigurable MRI coil, both aiming to reduce the coupling of implanted leads and MRI electric fields. Our results demonstrated that both techniques perform well under variant body compositions.

## I. Introduction

More than 12 million people in the united states are presently carrying a form of conductive medical implant such as cardiac pacemakers or neuromodulation devices [1]. It is estimated that 50-75% of these patients will need magnetic resonance imaging (MRI) exams during their lifetime [2], with many patients requiring repeated examination [3]. The major risk of performing MRI on patients with electronic implants is due to the radiofrequency (RF) heating of the tissue surrounding the implant’s tip, a phenomenon commonly known as the antenna effect [4]–[6]. In cases such as deep brain stimulation (DBS) devices where leads are implanted in sensitive organs such as the brain, highly restrictive guidelines are in place to safeguard patients, limiting MRI accessibility. DBS manufacturers for instance, have limited use of MRI to 1.5T horizontal scanners, and only with pulse sequences that generate specific absorption rate (SAR) of radiofrequency energy less than 0.1 W/kg (30 times below FDA limit for scanning in the absence of implants) or the magnetic field B_1_^+^rms ≤2μT [7]–[9]. Complying with these guidelines has proven to be difficult. For example, a typical stroke MRI protocol consists of T2/FLAIR, perfusion images and MR angiography [10]. These sequences have SAR and B_1_ levels well beyond what is allowed in patients with DBS implants. The situation is even more problematic within the context of musculoskeletal (MSK) imaging, for which the prescribed sequences have SAR levels already approaching the FDA limit for scanning even in the absence of implants. Among patients with movement disorders who are prone to falls and joint injuries, MSK MRI is often indicated, leaving those with DBS implants unable to receive the standard of care. Moreover, although currently 3T MRI is contraindicated for DBS patients, there are strong incentives to make it accessible for DBS imaging. 3T MRI confers a much better contrast-to-noise ratio compared to 1.5T MRI, allowing to delineate small structures surrounding DBS targets which is helpful for electrode localization [11].

Unsurprisingly, recent years have witnessed considerable efforts to reduce RF heating of DBS implants during MRI, both through MRI hardware modification, and by modifying the implanted lead’s design, material, and trajectory. From MRI hardware perspective, promising studies have shown the possibility of applying parallel transmit technology [12]–[20], and reconfigurable MRI coils [21]–[25] to shape the electric field of the MRI transmit coil in each individual patient such that field interactions with DBS leads is minimized. other efforts have focused on modifying the implant’s design and material [26]–[28] or its trajectory inside the body [29]–[31] to reduce its coupling with MRI electric fields. However, these studies have either relied on simulations in homogenous body models, or experiments with box-shaped single-material phantoms. This is a limitation, as the distribution of MRI electric fields inside a sample can be substantially altered by changes in local electric properties of the sample [32], which consequently affects RF heating of a conductive implant [33]. The problem is particularly important in the context of implant heating, as highlighted in our recent study with a commercial DBS device implanted in a multi-material anthropomorphic phantom reporting significant increase in RF heating in phantoms with subcutaneous fat compared to phantoms with no fatty tissue [34]. This raises the concern as whether or not the promising strategies previously introduced by reconfigurable MRI technology and surgical lead management are applicable in patients with diverse body characteristics.

In this work, we applied numerical simulations to investigate major concerns regarding applicability of RF coil modifications and DBS lead management strategies to reduce RF heating across patients with different body types. Specifically, we aim to address three open questions that will help advancing development of such techniques:

### 1) Can numerical simulations reliably predict alteration of RF heating due to variations in patient’s body composition?

As experiments to establish safety of implants in MRI environment are expensive and time consuming, there is an increasing trend towards application of numerical simulations for these purposes [35]–[39]. Recent reports on differences in the RF heating of DBS implants due to changes in phantom composition [34] warrant the need to investigate the electromagnetic mechanism underlying the phenomenon. We performed numerical simulations to investigate whether experimental observations in multi-material phantoms [34] could be replicated *in silico*, and whether more realistic body models with and without local fat around the implanted pulse generator (IPG) would show the same trend.

### 2) Are DBS lead management strategies that aim to reduce MRI heating effective in patients with diverse body types?

Modifying the extracranial trajectory of DBS leads can significantly reduce RF heating during MRI as shown in simulations with homogenous head models and experiments with single-material phantoms [29]–[31]. As presence of fat alters the distribution of MRI electric fields and subsequently changes the RF heating, it is important to examine whether or not lead management strategies will be effective across different body characteristics. In this work, we report simulation results with variant DBS lead configurations in body models with and without subcutaneous fat and examine the effectiveness of lead management techniques across different body types.

### 3) Are MRI hardware modification techniques based on field-shaping effective in patients with diverse body types?

In the context of MRI hardware modification for DBS imaging, reconfigurable patient-adjustable (RPA) coils have shown promise to reduce RF heating of DBS implants on a patient-specific basis [21]–[25]. The technique works by rotating a linearly-polarized (LP) birdcage transmit coil, which has a slab-like region of zero electric field, around patients head, such that implanted DBS leads are contained within the zero electric field region. To date, all studies that assessed feasibility and performance of RPA coils were performed with homogenous head models and homogenous phantoms. Here, we performed finite element simulations to quantify, for the first time, the degree to which SAR-reduction performance of an MRI reconfigurable coil is dependent on patient’s body characteristics.

This work aims to set the ground for further development and dissemination of techniques that reduce RF heating of DBS implants through MRI hardware modification and surgical lead management strategies. If successful, clinical implementation of such techniques would impact patient’s care in a large scale.

## II. The effect of subcutaneous fat on the RF heating of DBS devices during MRI with body coils

Recently we reported that patterns of RF heating of a DBS device implanted in an anthropomorphic phantom can be significantly altered due to changes in the phantom composition [34]. Here we investigate whether a similar pattern can be predicted by numerical simulations, and if such simulations can shed light on the electromagnetic mechanism underlying the phenomena.

### A. Experiments

Details of experimental setup are given in [34], and illustrated in Figure 1. In brief, an anthropomorphic phantom consisting of a human-shaped torso and skull was designed and fabricated along with grids and supporting structures that allowed DBS device to be positioned in a manner similar to the clinical practice. The skull was filled with saline-doped agar gel (*σ* = 0.40 *S/m* and *ε_r_* = 78) and inserted into the body of the phantom which was then filled with saline solution (*σ* = 0.50 *S/m* and *ε_r_* = 78) (Figure 1). A Medtronic DBS device (Medtronic Inc, Minneapolis, MN) with a 40 cm lead (model 3387), 60 cm extension (model 3708660) and an implantable pulse generator (IPG) (Activa PC-37601) was implanted in the phantom. MR compatible flouroptic temperature probes (OSENSA, BC, Canada) were secured to the lead contacts and the lead-probe system was inserted into the skull through a 5 mm hole drilled on the surface. The phantom was positioned in the MRI scanner such that its head was at the iso-center of the magnet, mimicking RF exposure during brain imaging. RF heating was measured for four different extracranial lead trajectories as shown in Figure 1. Experiments were repeated by replacing 7 L of saline by the same amount of vegetable oil which created a 3-cm layer of fat on top of the saline solution, mimicking presence of subcutaneous fat.

**Figure 1:**
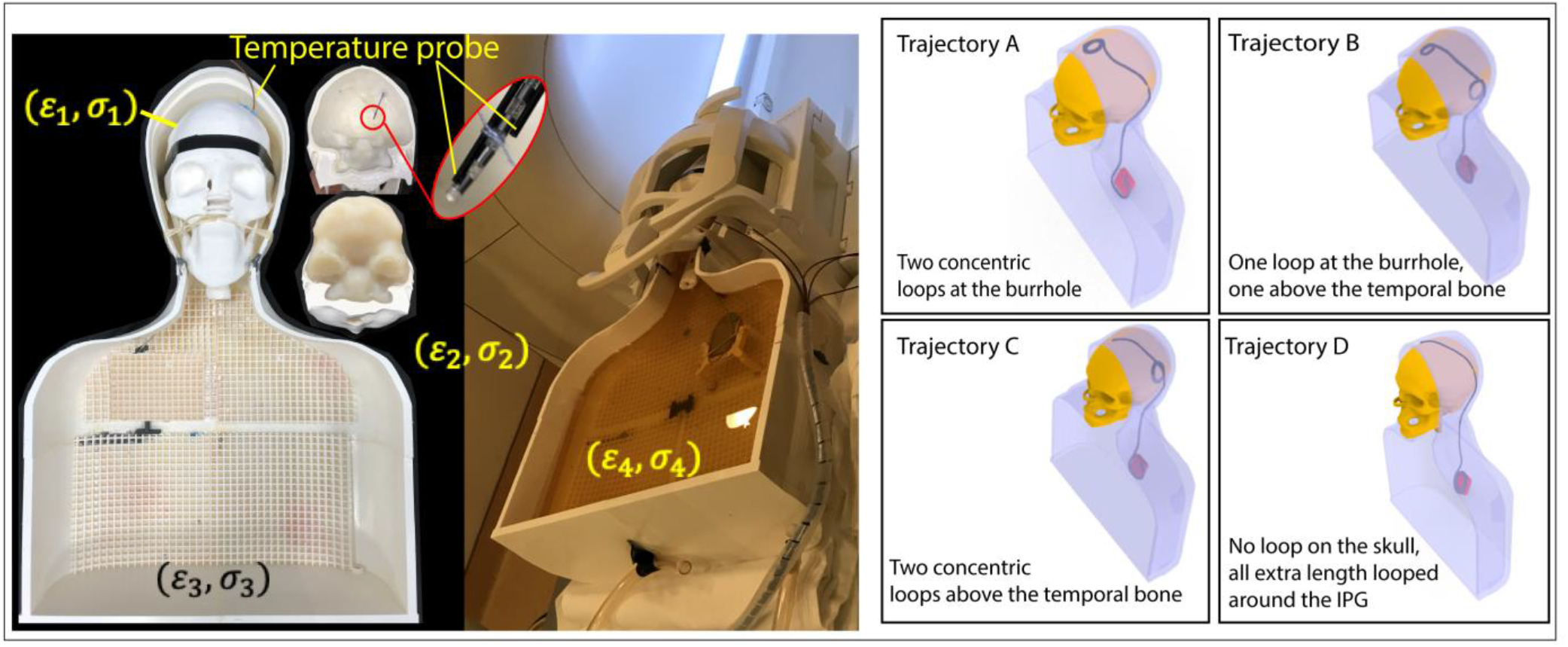
Left: Experimental setup showing the anthropomorphic phantom. The skull structure was 3D printed in plastic with low conductivity and permittivity (*σ* = 0 *S/m* and *ε_r_* = 3.5) similar to bone and was filled with brain mimicking gel (*σ* = 0.40 *S/m* and *ε_r_* = 78). The torso was filled with saline (*σ* = 0.50 *S/m* and *ε_r_* = 78) and oil (*σ* = 0 *S/m* and *ε_r_* = 3). The DBS device was positioned inside the phantom similar to clinical practice. Right: Different configurations of extracranial lead and extension trajectories, mimicking different surgical strategies.

RF heating measurements were performed at a Siemens 3T Prisma scanner (Siemens Healthineers, Erlangen, Germany), using body transmit coil and a 20-channel receive head coil. A T1 weighted turbo spin echo sequence with TE = 7.5 ms, TR = 1450 ms, flip angle = 150°, B_1_^+^rms = 2.8 μT and acquisition time of 7 minutes 31 seconds was used during RF exposure.

### B. Simulations

To investigate how variations in distribution of electromagnetic fields of the RF coil due to presence of fat contribute to the changes observed in RF heating, we replicated the experimental setup in finite element simulations. Simulations were implemented in ANSYS Electronic Desktop 2019 (ANSYS, Canonsburg, Pennsylvania, USA), following a combined finite element-circuit analysis approach [40] as implemented in our previous works [35]. The simulated phantom consisted of models of the brain, skull, and body similar to the experimental phantom, with all tissues assigned to the same dielectric properties as measured in the experiments and reported above. Four DBS lead trajectories mimicking experimental setup were created as illustrated in Figure 1. MRI RF coil was modeled as a 16-rug high-pass birdcage (67 cm length 61 cm diameter), tuned at 127 MHz (3 T), and driven in quadrature through two signal sources placed at the end ring on patient’s head side. The input power of the coil was adjusted such that it produced a mean *B*_1_^+^ = 2.8 μT on a circular plane at its iso-center, similar to the B_1_^+^ reported by the scanner during RF exposure experiments. The maximum of 1g-averaged SAR was calculated around the tip of DBS leads for each scenario to be compared with experimental results.

All simulations were performed twice, once with a phantom without the fat, and a second time with a layer of fat covering the surface of phantom’s body similar to the experimental setup. It is important to note however, that in reality, overweight patients have increased amount of local subcutaneous fat mostly in the chest area surrounding the IPG. This is different from our experimental setup, where the oil layer fully covered the surface of phantom’s body. To examine if a similar trend in variation of RF heating was present in a case that represented human body more realistically, we also performed simulations with a human-shaped body model consisting of average tissue (*σ* = 0.50 *S/m* and *ε_r_* = 78), skull (*σ* = 0.07 *S/m* and *ε_r_* = 15), brain tissue (*σ* = 0.40 *S/m* and *ε_r_* = 78), with and without a block of local fat (*σ* = 0.04 *S/m* and *ε_r_* = 6) covering the upper face of the IPG and initial segments of the extension as demonstrated in Figure 2B.

**Figure 2:**
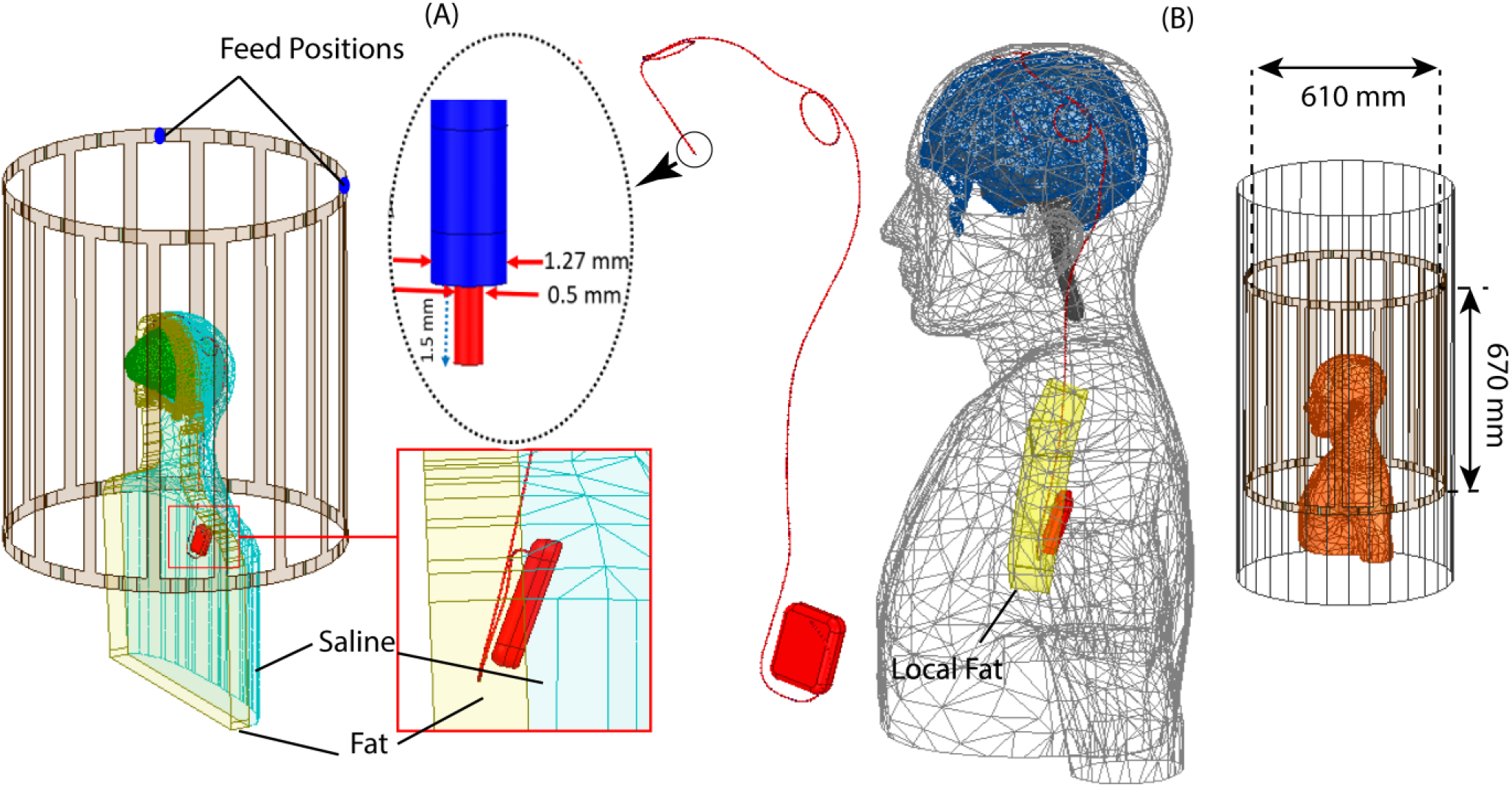
Simulation setups with experiment mimicking phantom model (A) and realistic body model (B), showing fat representing tissue for either case. The lead model with its tip characteristics also shown.

ANSYS HFSS follows an adaptive mesh scheme with successive refinement of an initial mesh between iterative passes. We set the initial mesh such that maximum element size was <2 mm for the lead, <4 mm for the insulation, <5 mm for IPG, <10 mm for the coil, <20 mm for brain, body, and coil shield, and <10 mm for local fat and skull. All simulations converged with 2-4 adaptive passes. Total time taken for each simulation was about two and half hours on a DELL server with 1.5 TB memory and 2_Xenon(R) Gold 6140 CPUs each having 32 processing cores. Table 1 shows mesh statistics for a typical simulation.

**Table 1:** Mesh statistics for a simulation with experiment mimicking phantom with trajectory D and no fat tissue.

| Parts | No of Tetrahedrons | Min. edge length (mm) | Max. edge Length (mm) | RMS edge length (mm) |
| --- | --- | --- | --- | --- |
| Lead | 44827 | 0.06 | 1.99 | 0.82 |
| Insulation | 85860 | 0.06 | 2.15 | 1.05 |
| CSF | 30716 | 0.11 | 27.05 | 11 |
| Skull | 79414 | 0.2 | 12.2 | 7.63 |
| SAR Box | 41704 | 0.04 | 2.41 | 1.57 |
| Body | 144681 | 0.1 | 25.17 | 14.47 |

### C. Results

Figure 3A shows the temperature rise Δ*T* measured at the tip of the DBS lead during RF exposure experiments with different lead trajectories of Figure 1 in phantoms with and without subcutaneous fat. Figure 3B gives the numerical results for the maximum 1gSAR in simulations that mimicked the experimental phantom setup. As it can be observed, the effect of variation in phantom composition and lead trajectories on Δ*T* is reflected in SAR simulations with a good agreement. Specifically, for the worst-case heating scenario (trajectory D), the presence of fat increased the measured Δ*T* by 7-fold (from 0.61 *°C* to 4.70°C) and the calculated 1gSAR by 6-fold (from 29.7 W/kg to 169.4 W/kg). Simulations with the realistic body model with and without local fat around the IPG predicted a similar pattern of change in the RF heating at the electrode tip: for the two lead trajectories with highest heating (trajectory D and C) the presence of local fat around the IPG increased the 1gSAR at the electrode tip by 140%, and 45% respectively.

**Figure 3:**
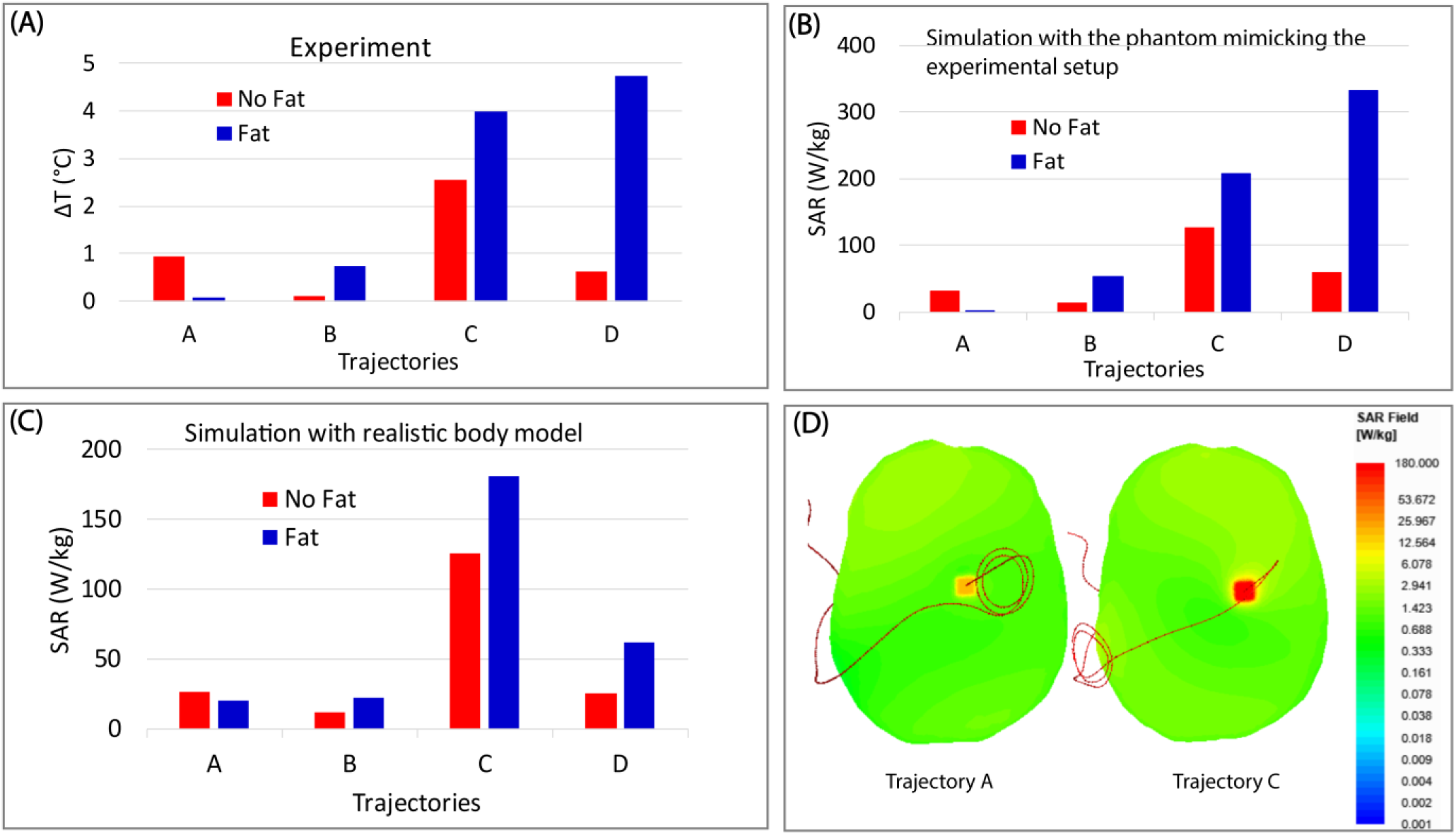
(A) Plot of experimentally measured temperature rise at DBS lead electrodes for lead trajectories A-D shown in Figure 1, in phantoms with and without subcutaneous fat. (B-C) The maximum of 1g-averaged SAR at the tip of the simulated DBS lead in a phantom that mimicked the experimental setup, and simulations with realistic body model with and without subcutaneous fat. (D) Plot of 1g-averaged SAR on an axial plane passing through the exposed tip of the lead for trajectories A and C in realistic body model with fat tissue.

Modifying lead trajectories to introduce concentric loops at the surgical burr hole (trajectories A and B) significantly reduced the RF heating compared to the trajectory without loop (trajectory D) or the trajectory with loops placed above the temporal bone (trajectory C). From Figure 3, it can be observed that implementing trajectory A, Δ*T* was reduced by 98% in the phantom with fat (from 4.7 0°C (Trajectory D) to 0.10°C) and by 63% in the phantom without fat (from 2.54°C (Trajectory C) to 0.93°C) compared to the corresponding worst case *ΔT.* The same trend was predicted by simulations, with respective reductions in SAR being 99% (compared to Trajectory D) and 76% (compared to Trajectory C) for simulations that mimicked our phantom experiments, and 89% (compared to Trajectory C)and 79% (compared to Trajectory C) for simulations with realistic body model.

RF heating of an implant is known to be highly affected by the background electric field of the MRI scanner [41]. To assess how the distribution of the background electric field was varied due to change in the phantom composition, we performed simulations in phantoms with and without fat in the absence of the implant. Figure 4A shows simulated maps of the magnitude of incident electric field on three coronal planes in the phantoms at 17 mm below, 3 mm above, and 23 mm above the fat-saline interface (the implant is shown only for visualization and was not included in simulations). The distribution of B_1_^+^ field is also given on the plane that was 3 mm above the fat-saline interface (plane 2) in Figure 4B. As it can be observed, the electric field magnitude is substantially higher on planes that pass through the fat layer, even though the B_1_^+^ field remained relatively unchanged. Consequently, lead trajectories that pass through fatty tissue will be exposed to a higher electric field resulting in higher induced currents, even though they experience same B_1_^+^ magnitude.

**Figure 4:**
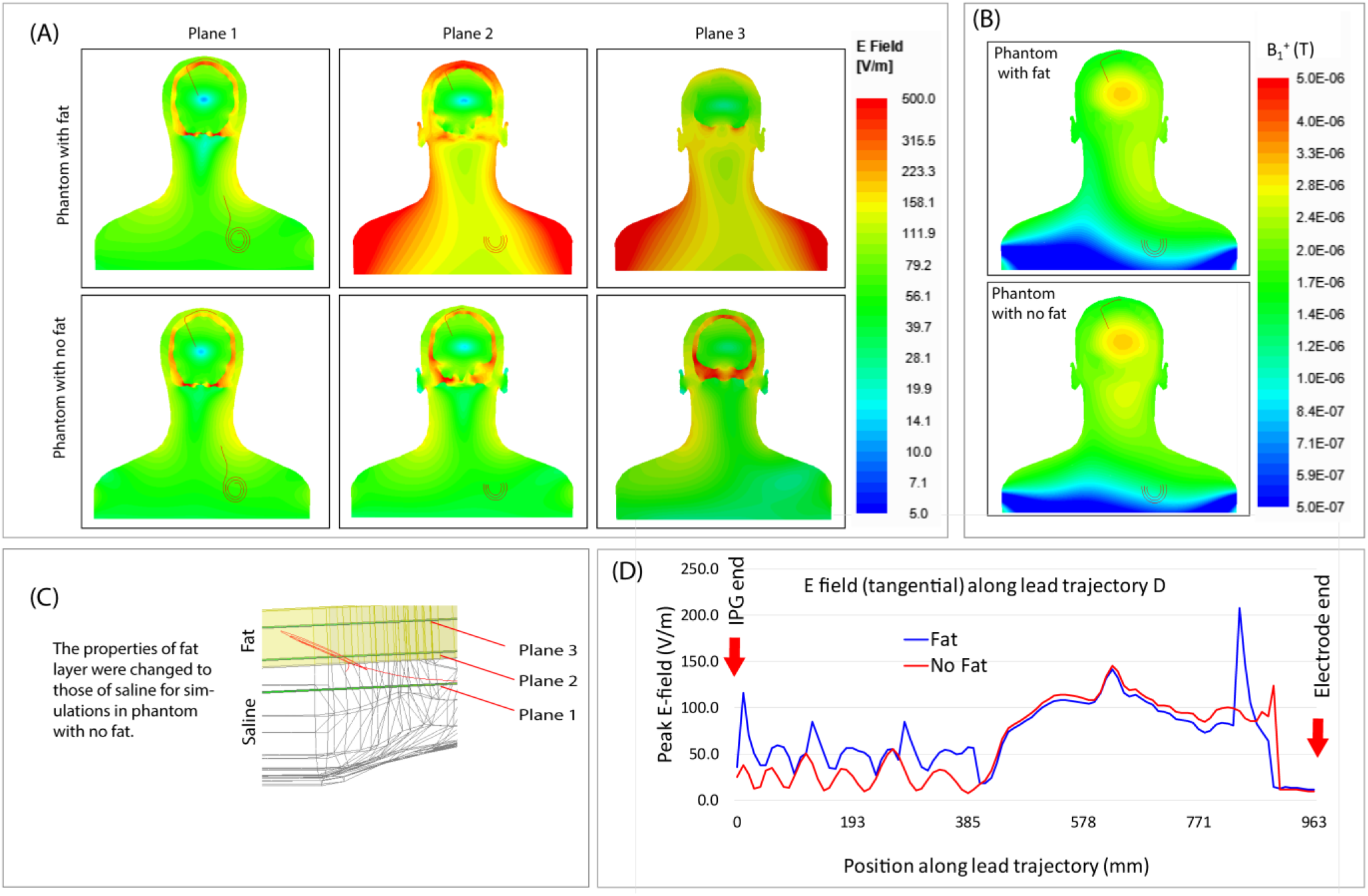
(A) Simulated maps of electric field (magnitude) distribution on three coronal planes positioned below and above fat-saline interface. Simulations are performed in phantoms with and without fat (where the properties of fat layer was changed to that of saline) in the absence of DBS implant. The implant is shown only for visualization. (B) Plot of B_1_^+^ on plane 2 in phantoms with and without fat. (D) Peak value of the tangential component of electric field along the length of the lead trajectory D, in phantoms with and without fat.

This can be better appreciated in Figure 4D where the peak value of the tangential component of the incident electric field, E_tan_, is shown along the lead trajectory D. On average, the magnitude of E_tan_ along the last 40 cm of the extension (the portion starting from the IPG) was 2.5 times higher in the phantom with additional fat than in phantom with only saline. Such increased incident electric field can lead to increased induced current along the DBS lead, resulting in increased RF heating, as theoretically predicted by the concept of lead transfer function [6], [42]–[44].

## III. Performance of RPA coil technology in patients with increased subcutaneous fat

The reconfigurable patient-adjustable (RPA) coil technology was recently introduced to reduce RF heating of DBS implants by rotating a linearly polarized birdcage transmit coil around patient’s head such that the implant could be contained within the low electric field region of the coil. This reduces induced electric currents on the leads which in turn reduces RF heating. To date, simulation studies and experimental measurements that aimed to characterize the performance of RPA technology have done so using homogenous head or body models [21]–[25]. Here we investigated the performance of a 3T RPA coil in reducing RF heating of fully implanted DBS systems, and examined if increased amount of subcutaneous fat around the IPG would affect the SAR-reduction efficiency of the coil.

### A. Simulations

We performed finite element simulations with a model of a shielded high-pass birdcage head transmit coil (16-rung, 23.6 cm length, 30.6 cm diameter), tuned to 127 MHz (3 T), and driven in linear mode by a single sinusoidal source placed at one of the end rings. The coil was rotated around the head of the heterogeneous body model of Figure 2 with 11.25° increments to cover a full circle (Figure 5). The maximum of 1g-averaged SAR was calculated around the tip of the leads for trajectories A-D implanted in body models with and without local fat around the IPG. Simulations were repeated by replacing the linearly polarized birdcage head coil with a birdcage body coil driven in quadrature mode (circular polarization) for calculation of SAR reduction efficiency (SRE). For all simulations, the input power of the body and the RPA coils were adjusted to produce mean B_1_^+^ =2 μT on an axial plane passing through center of the coil.

**Figure 5:**
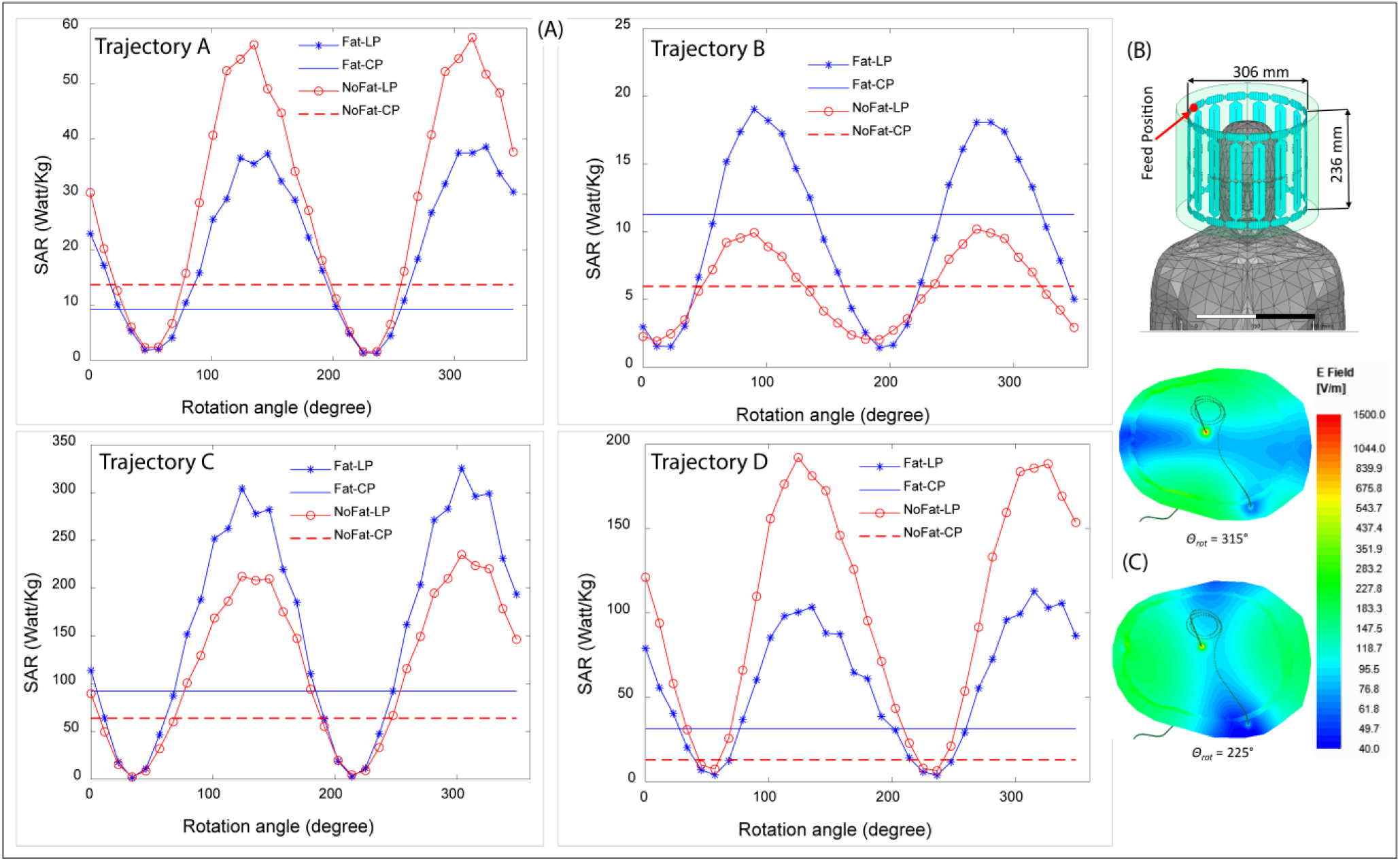
(A) Plot of maximum value of SAR (1g averaged) against angle of rotation of RPA coil for all four different trajectories in the presence and absence of fat tissue. The horizontal lines represent the SAR values generated by the CP body coil. All the SAR values are normalized to B_1_^+^ =2μT. (B) RPA coil with body model inserted. The position of feed position corresponding to θ = 0° is shown in red. The coil was rotated around patient’s head with 11.25° increments. (C) Complex magnitude of E-field on a transverse slice, showing changes in orientation of low E-field band with rotation of the RPA coil.

The maximum SRE of the RPA coil was quantified as:

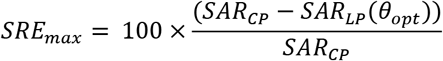

where *SAR_CP_* is the maximum of 1g-averaged SAR around the DBS lead generated by the CP body coil, *θ_opt_* is the optimum angle of the RPA coil that maximally reduces the SAR, and *SAR_LP_*(*θ_opt_*) is the maximum of 1g-averaged SAR when the RPA coil is positioned at its optimal angle.

### B. Results

Figure 5A gives the maximum of 1g-averaged SAR at the tip of the lead as a function of RPA coil’ s angle for different lead trajectories implanted in body models with and without local fat around the IPG. The maximum of 1g-averaged SAR generated by the CP body coil is also given for comparison. As it can be observed, for all four trajectories the SAR generated by the RPA coil at its optimum rotation angle was significantly lower than the SAR generated by the body coil, irrespective of the presence or absence of fatty tissue. Additionally, the optimum angle for SAR reduction remained unchanged by the inclusion of fatty tissue for all trajectories. Figure 5C gives the distribution of E field for two positions of the RPA coil corresponding to maximum and minimum SAR.

The SAR reduction efficiency of the RPA coil for different lead trajectories implanted in phantoms with and without fatty tissue are given in Table 2. The maximum *SRE =* 97.1% was achieved for the lead with trajectory C implanted in the body model with fat. The minimum SAR reduction was observed for lead trajectory D implanted in the body model without fat tissue with *SRE* = 49.6%. For trajectories producing lower heating (Trajectories A and B), the SAR reduction was more than 65% for each case with or without fat tissue.

**Table 2:**
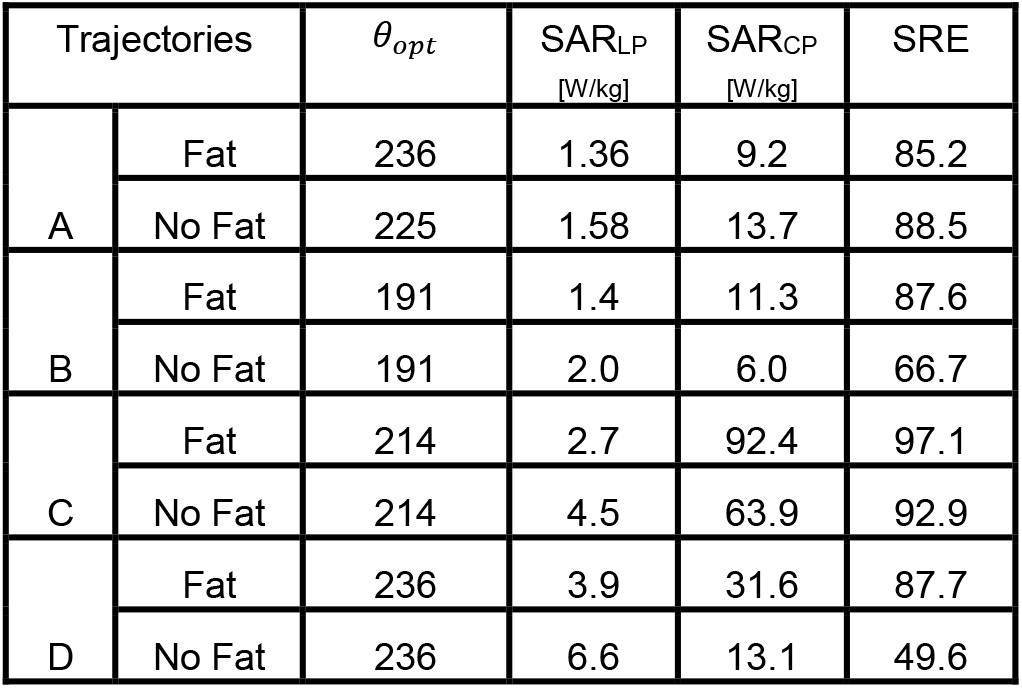
SAR reduction efficiencies (SRE) at optimal angle of rotation for each trajectory for different body composition.

| Trajectories | | $\theta_{opt}$ | SAR<sub>LP</sub><br>[W/kg] | SAR<sub>CP</sub><br>[W/kg] | SRE |
| --- | --- | --- | --- | --- | --- |
| A | Fat | 236 | 1.36 | 9.2 | 85.2 |
|  | No Fat | 225 | 1.58 | 13.7 | 88.5 |
| B | Fat | 191 | 1.4 | 11.3 | 87.6 |
|  | No Fat | 191 | 2.0 | 6.0 | 66.7 |
| C | Fat | 214 | 2.7 | 92.4 | 97.1 |
|  | No Fat | 214 | 4.5 | 63.9 | 92.9 |
| D | Fat | 236 | 3.9 | 31.6 | 87.7 |
|  | No Fat | 236 | 6.6 | 13.1 | 49.6 |

Finally, we examined the performance of the RPA coil in terms of B_1_^+^ inhomogeneity by calculating the ratio of the standard deviation and mean of B_1_^+^ on a transverse plane inside the head located 2cm above the coil’s iso-center (Figure 6 (A)) as:

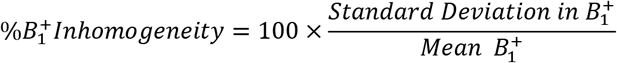

**Figure 6:**
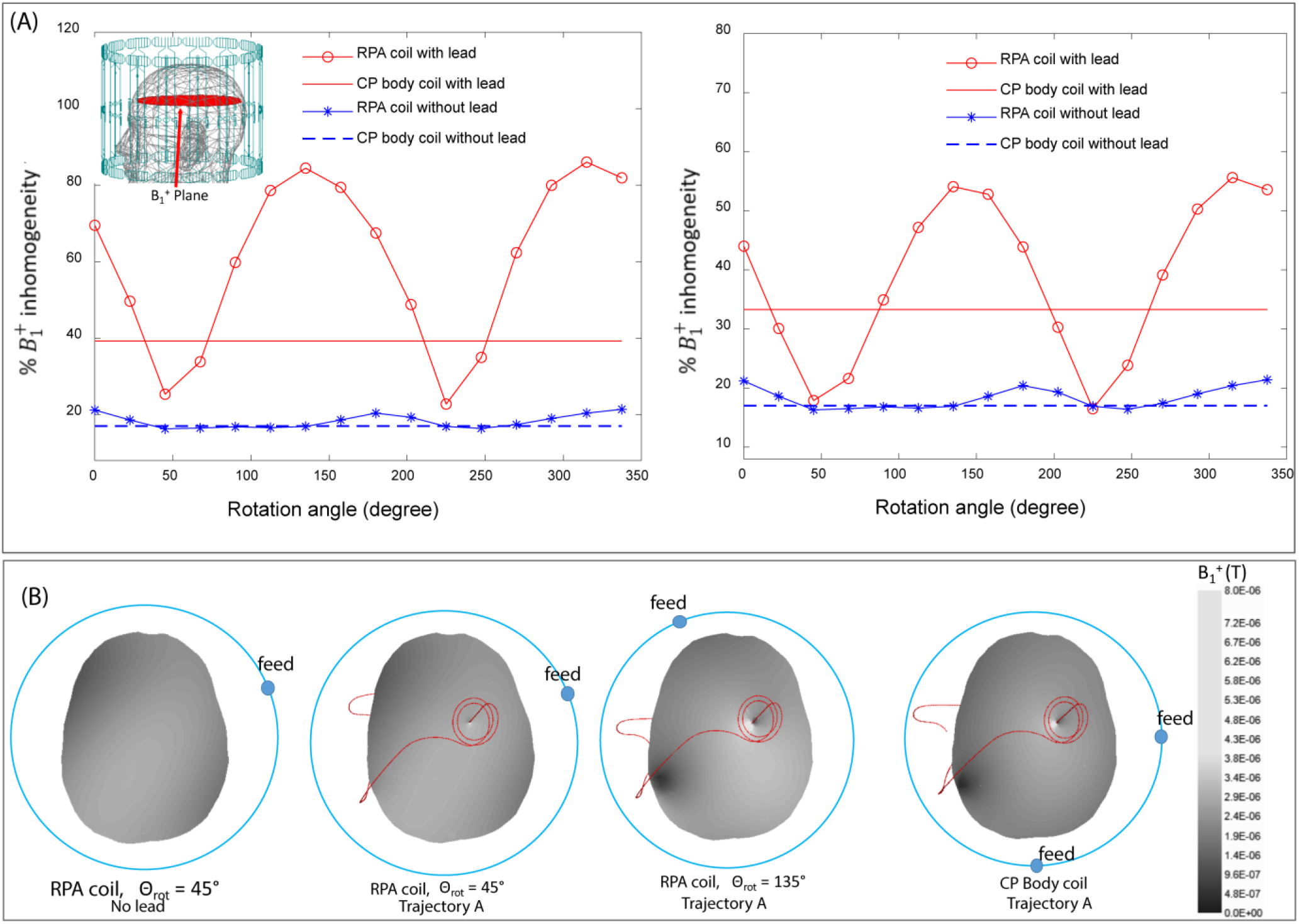
(A) Plot of B_1_^+^variation with angle of rotation of RPA coil, in the presence DBS lead (trajectory A) as well as without lead using realistic body model without fat tissue (left) and with fat tissue (right). The corresponding variations for CP body coil are also added as horizontal lines. The transverse plane was covering brain tissue only. (B) Plot of B_1_^+^ field distribution in the same plane for RPA coil as well as CP body coil. For RPA coil, the plots correspond to minimum B_1_^+^ inhomogeneity (*θ_rot_* = 45°) and maximum B_1_^+^ inhomogeneity (*θ_rot_* = 135°). The position of feed in each case has been shown by the dot on the circle (figures not to the scale).

Figure 6 (A) shows the plots of B_1_^+^ inhomogeneity for lead trajectory A, implanted in the realistic body model with and without fat tissue using both RPA coil and the CP body coil. Reference values for B_1_^+^ inhomogeneity in the absence of the implant are also included for comparison. It has been shown that the B_1_^+^ field of an empty linearly polarized (LP) birdcage coil is highly uniform, however, this changes when the coil is loaded with a human head [45]. Field inhomogeneity inside the head will be specifically affected by the location of the linear feed with respect to the head. From Figure 6, the maximum B_1_^+^ inhomogeneity in the head in the absence of the implant was 21.4% for the RPA coil (compared to 17% for CP body coil), which occurred when the linear feed was located 45° behind the ear. The field inhomogeneity was, however, reduced to 16.3% when the linear feed was located between nose and right ear. Interestingly, this is the position that also reduces the RF heating of most typical DBS leads [25].

Presence of metallic implants can worsen the B_1_^+^ field inhomogeneity, as the induced RF currents along the implant generate secondary magnetic fields, thus distorting the original field distribution [14], [46], [47]. Plots in Figure 6 (A) clearly show that the B_1_^+^ field inhomogeneity variation with rotation of RPA coil, in the presence of implant, follows the same trend as the RF heating, with minima and maxima positions coinciding with the SAR minima and maxima (Figure 5). Additionally, at optimal position, the B_1_^+^ field inhomogeneity for RPA coil is reduced below the corresponding level for CP body coil, with as much as 35% improvement in field inhomogeneity in the phantom without fat and 46% improvement in the phantom with fat. From Figure 6B it is observed that B_1_^+^ field distortion due to presence of implant is minimized when the RPA coil is at its optimal position. For higher heating cases, significant B_1_^+^ distortion is produced not only by the intracranial part of the DBS lead, but also by the extracranial portions of the lead and extension. The results are consistent with earlier studies involving variation of RF heating and B_1_^+^ field inhomogeneity/image artifacts [14], [48].

## IV. Discussion and Conclusion

Patients with DBS devices can reap much benefit from MRI if their eligibility for MRI is extended to a wider range of sequences at 1.5 T as well as to higher field strengths beyond the restrictions imposed by current guidelines. Accessibility to 3 T MRI which is currently contraindicated for DBS can help provide better images of subcortical structures producing enhanced contrast compared to 1.5 T MRI [11]. This will lead to improvement in overall treatment procedure by reducing errors in target verification as well as enabling post-operative monitoring of induced changes in the function of affected brain networks. As the primary concern for contraindication is risks associated with RF heating, techniques for mitigating such heating can pave the way towards allowing use of 3 T MRI for DBS imaging. Due to high degree of complexity of RF heating phenomena [5], [49]–[53], the results of studies using simplified homogenous models might not provide enough confidence on patient’s safety and hence, more contributions incorporating complexity of body tissues are required. This study, using experimental measurements as well as established numerical simulation, has provided evidence that patient’s body composition has significant effect in RF heating of DBS devices during MRI. A more important upshot of this work, however, is that it affirms that risk mitigation strategies based on reconfigurable coil technology and DBS surgical lead management are still effective during 3 T MRI, even with variant patient’s body compositions.

In the past few years, numerical simulations have been increasingly used to assess safety of medical devices and imaging instruments [36], [54]–[58]. An important aspect of such practice is to validate simulations against measurements whenever possible, in order to provide confidence in future predictions of such models. Here we show that numerical simulations agree well with experiments in predicting the effect of modification in device configuration as well as patient’s body composition on RF heating of DBS implants. Addition of fatty tissue substantially altered the RF heating in simulations, which was in line with what was observed in our experimental setup. Simulations with body models that more realistically resembled human subjects predicted similar trend in variation in RF heating.

The alteration of RF heating at the tip of a DBS implant due to presence of local fat around the IPG might be due to two possible effects. Firstly, the local distribution of incident electric field of MRI scanner will be altered in and in close vicinity of the fat which has low permittivity and conductivity compared to surrounding tissue [32], [59]. This can in turn change the coupling of electric field with the portion of the lead/extension that is inside or close to the fatty tissue. Secondly, the presence of low permittivity fat will change the wavelength of RF fields surrounding the leads which can result in shifts in antenna resonance lengths [49], [50]. The substantial alteration of heating patterns due to presence of fat highlights the importance of considering body complexity in RF heating evaluations for implants during MRI.

Another important observation, confirmed with both simulations and measurements, was that lead trajectories with loops positioned at the surgical burr hole, or loops positioned both at the burr hole and toward the temporal bone, maximally reduced the RF heating for both cases with and without inclusion of fat. This suggests that lead management strategies are resilient to variations in patient’s body characteristics. It is worthy to note, however, the trajectories with loops placed at the burr hole are easier to implement surgically than the others.

In the context of MRI hardware modification, the performance of the RPA coil technology has not been tested on fully implanted DBS systems using patient models with variant body compositions. Here we showed that when positioned at its optimal angle, RPA coil consistently reduced RF heating compared to the CP body coil for all trajectories irrespective of inclusion or exclusion of fatty tissue. More importantly, the optimal angle for a particular trajectory was relatively unchanged for different body compositions. These finding indicate that effect of patient’s characteristics on the optimal rotation angle of the coil is small or negligible, although more study is required to draw definitive conclusions on the degree of coil sensitivity to body characteristics. The RPA coil comes with additional benefit of reducing B_1_^+^ inhomogeneity which is generated by induced current in implanted leads. When positioned at optimal rotation angle for SAR minimization, the induced currents along the lead are also minimized resulting into concurrent reduction in the B_1_^+^ field distortion. Such inhomogeneity is the source of the well-known nonsusceptibility image artifact around elongated implants [46].

Our results show that once the RPA coil technology is used together with device trajectory modification, its SAR reduction efficiency is significantly enhanced. Employing such techniques can substantially reduce the risk of RF heating in DBS patients during MRI at higher fields. This can open a way for wider accessibility of MRI in DBS patients beyond the limitations imposed by current guidelines leading to improvements as well as advancements in overall DBS treatment procedure.

Though this study includes tissue heterogeneity by including brain, skull and fat, it does not account for the heterogeneity inside the brain tissue, which can be a subject of further evaluation. Furthermore, representation of fatty tissue in this study has been quite simplified, using either a surface layer or a block of tissue around IPG. Using virtual population models that incorporate a wide distribution of fat can provide additional insight in the future. In addition, the results presented here are based on one model of DBS device (Medtronic, lead 3387, extension 3708660). Further study is required to draw conclusions for a wider range of DBS implant models being used in current practices.

## Acknowledgment

This work was supported by NIH grant R00EB021320.

